# Proportional variability relative to lower abundances: An interpretable and efficient index of deep lows in population time series

**DOI:** 10.1101/2022.02.19.481101

**Authors:** Mario Schlemmer

**Affiliations:** University of Klagenfurt

**Keywords:** ecological stability, extinction risk, proportional variability, lower-abundance baseline, computational efficiency, population crashes

## Abstract

Variability in population abundances is usually measured with the coefficient of variation (CV) or the standard deviation of log-transformed data (SDL), both of which are biased for non-Gaussian dynamics and zero counts. Proportional variability (PV_hi_), which scales each pairwise difference relative to the larger of the two abundances, avoids reference to the mean but responds only weakly to isolated crashes and to the depth of declines. I introduce PV_lo_, an index of proportional variability relative to lower abundances, based on all pairwise comparisons and bounded between 0 and 1. Zero counts are handled without data transformation. A sorting-based algorithm reduces the computation from *O*(*n*^2^) to *O*(*n* log *n*), making the index applicable to long time series. In example series and simulations (Gaussian, Cauchy, rare-crash, bimodal and gradual-decline regimes), PV_lo_ remained sensitive to crashes and to the contrast between abundance levels, whereas PV_hi_ levelled off, but PV_hi_ was more robust against heavy-tailed extremes. In a comparison with series of ordinary fluctuations, PV_lo_ and SDL separated crashes better than PV_hi_ and the CV only if the ordinary variability of a population was unknown, and PV_lo_ did not outperform SDL; during a sustained decline the indices differed little. PV_lo_ is a computationally efficient index that responds to deep lows on an interpretable ratio scale without transformation of the data, an alternative to SDL that is relevant for screening many populations in a conservation context.

## 1 Introduction

The temporal variability of population abundances is a central quantity in population ecology and conservation biology. It is used to compare the dynamics of populations and species and to assess extinction risk, because populations that fluctuate strongly are more likely to reach abundances at which demographic stochasticity and random catastrophes can eliminate them (Pimm, 1991; Lande, 1993). The IUCN Red List criteria, for example, list extreme fluctuations, typically more than one order of magnitude, among the conditions used to assess extinction risk (IUCN Standards and Petitions Committee, 2024). The monitoring data on which such comparisons rely are often short, contain zero counts and are punctuated by rare events such as crashes after extreme weather, outbreaks or local extinctions. These are the conditions under which traditional indices behave poorly (McArdle et al., 1990; Gaston & McArdle, 1994; Heath, 2006). Traditionally, variability in population abundances is quantified with ad hoc measures that express dispersion relative to an average. The coefficient of variation (CV) divides the standard deviation by the mean of the original data; the standard deviation of log-transformed abundances (SDL) applies the same logic on the logarithmic scale. Both measures have well-known shortcomings. The CV is dominated by single large values and cannot separate populations that differ in the depth of their lows, and its behaviour is closely tied to the mean abundance (McArdle et al., 1990; Heath, 2006). SDL is better suited to multiplicative dynamics, but it is undefined for zero counts, which are common in monitoring data. The usual remedy, adding a constant (typically 1) to all abundances before taking logarithms, is arbitrary: the scores depend on the size of the constant relative to the abundances and hence on the unit of measurement (Gaston & McArdle, 1994). In addition, SDL is expressed in logarithmic units and has to be back-transformed to be interpreted in terms of abundances. Neither measure has a constant range, despite being relative, so that a given score has no fixed meaning. Results from both have been described as unreliable when the underlying dynamics are not Gaussian, and their dependence on the mean has been identified as the chief reason (Heath, 2006). To quantify variability on a truly proportionate scale without reference to the mean, Heath (2006) introduced a framework that estimates variability from comparisons between all unique pairs of abundances. In its original formulation, this measure scales each pairwise difference exclusively relative to the higher value of the pair. To contrast this approach explicitly with the new measure that scales relative to the lower value of each pair, I refer to Heath’s (2006) original Proportional Variability index as PV_hi_ throughout this paper. It is defined as:

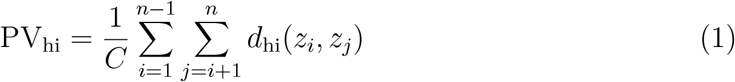

where the pairwise directional differential *d*_hi_ is given by:

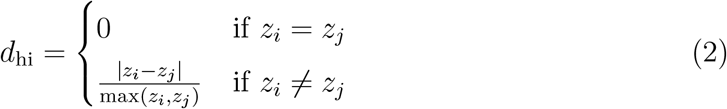

The total number of pairs of observations is *C* = *n*(*n*−1)*/*2, where *n* is the length of the time series. PV_hi_ has clear merits: it is bounded between 0 and 1, it does not depend on the mean, and zero counts enter the comparisons without any transformation (*d*_hi_ = 1 for every pair that contains a zero). Its weakness is that it averages proportional declines, which are themselves bounded at 100%. It therefore makes little difference whether a population has lost 90% or 99% of its abundance (*d*_hi_ = 0.90 versus 0.99), although the second case means a hundred-fold and the first a ten-fold difference in abundance. As a consequence, PV_hi_ levels off once abundances differ by about an order of magnitude, and isolated crashes, however deep, add only a small and nearly constant amount to the average. For extinction-risk assessment, however, it is the depth of the lows that matters, because demographic stochasticity and random catastrophes act mainly on small populations (Pimm, 1991; Lande, 1993). The commonly used measures therefore leave a gap: CV is dominated by high values, PV_hi_ is only weakly sensitive to low ones, and SDL, although sensitive to low abundances, requires an arbitrary constant when zeros occur and yields values without direct meaning on the scale of the abundances.

Here I introduce PV_lo_, which scales each pair relative to the lower abundance, so that the same two cases differ by 9 and 99 times the lower abundance. The index rests on four properties. First, it is sensitive to the depth of lows, and it keeps increasing when abundances differ by more than one order of magnitude. Second, it can be read on a ratio scale: 1 + pv_lo_ is the average ratio of the larger to the smaller abundance of a pair, and the standardized index PV_lo_ is bounded between 0 and 1, so that a score can be interpreted directly without back-transformation. Third, zero counts are handled without transforming the data: no constant is added to the abundances, and only pairs that contain a zero are affected by the convention of counting in single individuals. Fourth, a sorting-based algorithm reduces the computation from *O*(*n*^2^) to *O*(*n* log *n*), which removes the main practical disadvantage of pairwise indices and makes the screening of many long series feasible. These properties do not make PV_lo_ a universally superior measure. It is less robust than PV_hi_ against heavy-tailed extremes, and, like SDL, which it resembles in its response to low abundances, it ignores the temporal order of the observations. It is designed for a specific question, namely how far abundances fall below the levels that a population usually attains, and for this question it offers SDL-like sensitivity on a scale that is directly interpretable. I describe the algorithm and compare PV_lo_ with CV, SDL and PV_hi_ for constructed example series and for simulated time series, following the simulation design of Heath (2006) and supplemented by a gradual-decline regime and a comparison of crash series with ordinary fluctuations.

## 2 Methodology

### 2.1 Proportional Variability Relative to Lower Abundances

Based on all pairwise comparisons of abundances in a time series, the average proportional difference relative to lower abundances is defined as

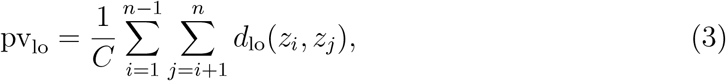

where *z*_*i*_ and *z*_*j*_ are the abundances of the *i*-th and *j*-th observation, and the pairwise differential *d*_lo_ is given by

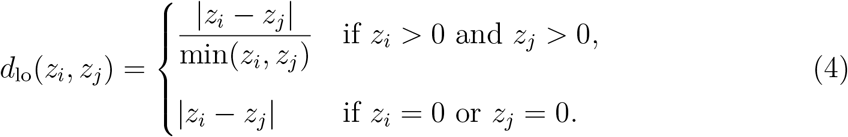

If a pair contains a zero, the difference is divided by 1 (a single individual) instead of by the lower value 0, which avoids a division by zero.

The score pv_lo_ has no upper bound. To constrain it to the unit interval while preserving its ordering, a standardized index is defined as

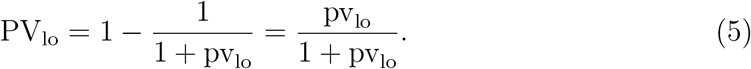

For two positive abundances, 1 + *d*_lo_ is the ratio of the larger to the smaller value, so 1 + pv_lo_ is the average of these ratios and PV_lo_ can be read directly on the original scale. A score of PV_lo_ = 0.25 means that the difference between two observations is on average one third of the smaller value (average ratio 1.33). At PV_lo_ = 0.50 the average difference equals the smaller value (average ratio 2), and at PV_lo_ = 0.90 it is nine times the smaller value (average ratio 10). For a pair containing a zero, 1 + *d*_lo_ = (*z* + 1)*/*1, i.e. the ratio is formed after adding 1 to both members of that pair. Since all pairs are compared irrespective of their temporal order, high scores indicate that low abundances lie far below high ones; they do not distinguish crashes from recoveries.

The index is directly related to Heath’s (2006) PV_hi_. For every pair of positive abundances, *d*_lo_*/*(1 + *d*_lo_) = *d*_hi_. With *f* (*x*) = *x/*(1 + *x*) one therefore has, for series without zeros, 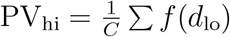, the average of the standardized pairwise differences, whereas 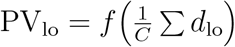 standardizes the average difference. Because *f* is concave, Jensen’s inequality gives PV_lo_ ≥ PV_hi_ in a series without zero counts, with equality only if all pairwise differentials are equal (a constant series, or *n* = 2). For pairs containing a zero the identity does not hold (Section 2.2).

### 2.2 An Efficient Computational Approach

The standard ad hoc measures of variability, such as CV and SDL, offer the clear advantage of being computationally efficient, even for exceptionally long time series. In contrast, truly proportional metrics have traditionally required the calculation of all *C* = *n*(*n* − 1)*/*2 unique pairs of abundances (Heath, 2006), a task whose cost scales quadratically (*O*(*n*^2^)) and becomes prohibitive for long or high-frequency monitoring series.

The pairwise sums can, however, be evaluated without enumerating the pairs. Since none of the indices depends on temporal order, the series can be relabeled: from here on, *Z* = (*z*_1_, *z*_2_, …, *z*_*n*_) denotes the series sorted in ascending order, *z*_1_ ≤ *z*_2_ ≤ *· · ·* ≤ *z*_*n*_ (ties in arbitrary order), and the index *i* refers to the rank. Let 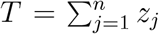 be the total abundance. For the *i*-th value in this order define the sums of all values ranked below and above it,

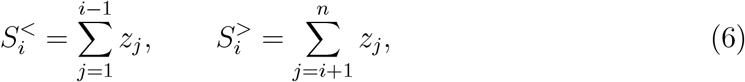

and the numbers of values ranked below and above it, 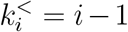 and 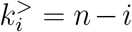. With the cumulative sums *c*_*i*_ = ∑_*j*≤*i*_ *z*_*j*_ one has 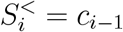 and 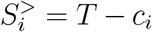, so that all sums are available in *O*(*n*) time after sorting and the total effort is dominated by the sort, *O*(*n* log *n*). The principle is the one used to compute Gini’s mean difference from the ordered sample without forming all pairs (David, 1968; Yitzhaki & Schechtman, 2012); here it is applied to pairwise ratios instead of differences, which is possible because the denominator of each pair is fixed by its rank. Counting ranks instead of values makes the procedure valid for tied abundances: a tied pair contributes (*z* − *z*)*/z* = 0 to both indices, whichever member is ranked first.

Let pd_hi,*i*_ be the sum of *d*_hi_ over all pairs in which *z*_*i*_ is the larger member, and pd_lo,*i*_ the sum of *d*_lo_ over all pairs in which *z*_*i*_ is the smaller member. Because every pair (*j, i*) with *j < i* appears exactly once in pd_hi,*i*_ and once in pd_lo,*j*_, summing over *i* covers each pair once. For strictly positive abundances,

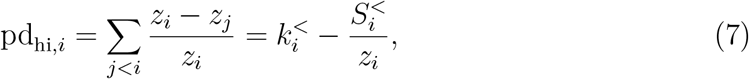

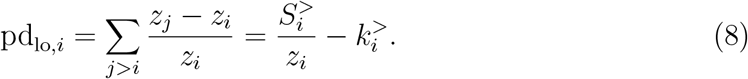

Since 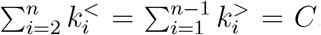, the indices follow directly from the cumulative sums. Heath’s index, which is the average of the pairwise differentials *d*_hi_ and is therefore already bounded between 0 and 1, is

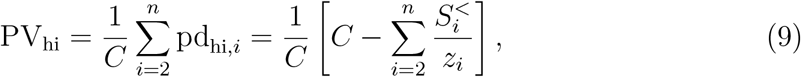

and, correspondingly, the non-standardized lower-proportional variability is

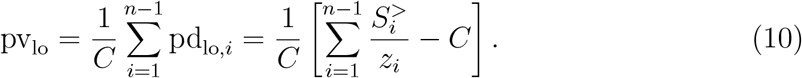

PV_lo_ is then obtained from pv_lo_ by the standardization in Eq. (5). This shortcut allows the proportional variability indices to be applied to high-frequency or long-term ecological monitoring data without compromising computational performance.

Figure 1 shows the Lorenz curve of the ordered abundances (Lorenz, 1905). Let *x*_*i*_ = *i/n* and *y*_*i*_ = *c*_*i*_*/T* be the coordinates of the Lorenz curve. The right-angled triangle below the *i*-th segment has legs 1*/n* and *z*_*i*_*/T* and thus the area *z*_*i*_*/*(2*nT*), i.e. it represents the abundance *z*_*i*_. In panel (a) the triangle above the segment has its apex in the origin and the area 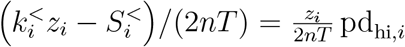; in panel (b) its apex is in the end point (1, 1) and the area is 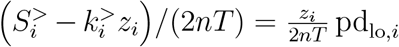. Each ratio of the area of the triangle above a segment to the area of the triangle below it therefore equals pd_hi,*i*_ or pd_lo,*i*_, respectively. The calculation of these proportional differences for each abundance is demonstrated in Table 1 using a worked example.

**Table 1:**
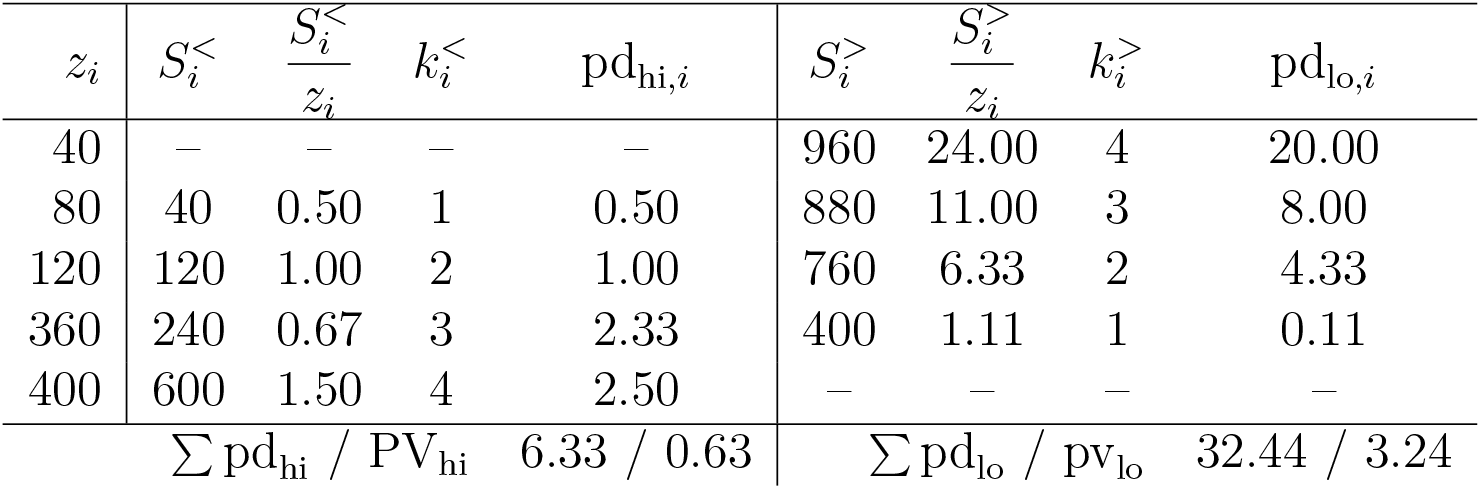
Step-by-step evaluation of the directional differentials pd_hi_ and pd_lo_ for the ordered abundance vector *Z* = (40, 80, 120, 360, 400) with *C* = 10.

| z<sub>i</sub> | S<sub>i</sub><sup><</sup> | S<sub>i</sub><sup><</sup>/z<sub>i</sub> | k<sub>i</sub><sup><</sup> | pd<sub>hi,i</sub> | S<sub>i</sub><sup>></sup> | S<sub>i</sub><sup>></sup>/z<sub>i</sub> | k<sub>i</sub><sup>></sup> | pd<sub>lo,i</sub> |
|---|---|---|---|---|---|---|---|---|
| 40 | – | – | – | – | 960 | 24.00 | 4 | 20.00 |
| 80 | 40 | 0.50 | 1 | 0.50 | 880 | 11.00 | 3 | 8.00 |
| 120 | 120 | 1.00 | 2 | 1.00 | 760 | 6.33 | 2 | 4.33 |
| 360 | 240 | 0.67 | 3 | 2.33 | 400 | 1.11 | 1 | 0.11 |
| 400 | 600 | 1.50 | 4 | 2.50 | – | – | – | – |
| ∑ pd<sub>hi</sub> / PV<sub>hi</sub> |  |  |  | 6.33 / 0.63 | ∑ pd<sub>lo</sub> / pv<sub>lo</sub> |  |  | 32.44 / 3.24 |

**Figure 1:**
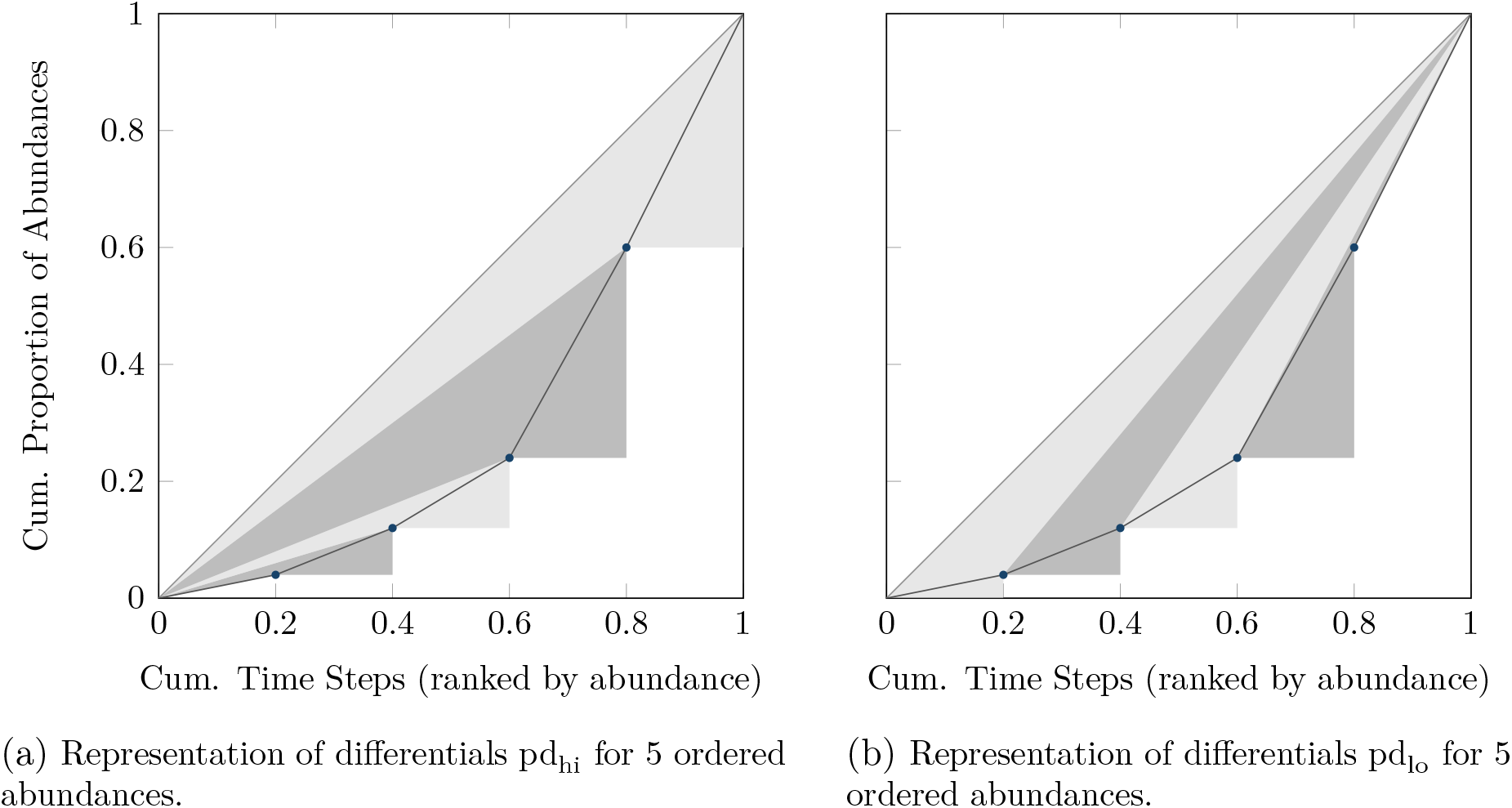
The Lorenz curve of the abundance vector (40, 80, 120, 360, 400). The area of the right-angled triangle below a segment is proportional to the abundance *z*_*i*_; the ratio of the area of the triangle above the segment (same shading) to the area of the triangle below it equals the differential pd_hi,*i*_ in (a) and pd_lo,*i*_ in (b) (Table 1). The triangle for *z*_1_ in (a) and for *z*_*n*_ in (b) is omitted because the corresponding differential is zero.

The efficient method extends directly to time series containing zero counts. The sums in Eqs. (9) and (10) divide by *z*_*i*_ and therefore require strictly positive abundances. When a time series contains zeros, all quantities in these equations (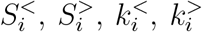 and the pair count) are therefore computed over the positive abundances only, and a correction term is added for the pairs that combine a zero with a positive value. Let *N*_0_ and *N*_+_ denote the numbers of zero and positive observations (*n* = *N*_0_ + *N*_+_), *C*_+_ = *N*_+_(*N*_+_ − 1)*/*2 the number of pairs of positive observations, and *T* the total abundance (which equals the sum over the positive observations). The ranks *i* in the following sums refer to the sorted positive observations. Then

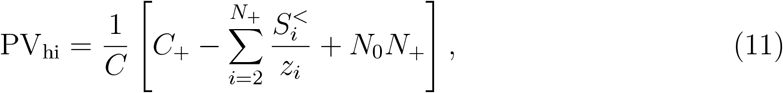

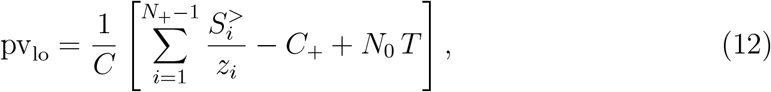

where the normalization still uses the pair count of the complete series, *C* = *n*(*n*−1)*/*2. For *N*_0_ = 0 these expressions reduce to Eqs. (9) and (10). Pairs of two zeros are included in *C*, but contribute 0 to both *d*_hi_ and *d*_lo_ and therefore do not appear in the correction terms; these cover only the *N*_0_*N*_+_ pairs that combine a zero with a positive value. As a check, the series *y*_9_ = (0, 0, 0, 0, 0, 0, 0, 0, 1, 99) in Table 2 has *N*_0_ = 8, *N*_+_ = 2, *T* = 100, *C*_+_ = 1, *C* = 45; the positive part contributes 99*/*1 − 1 = 98, and Eq. (12) gives pv_lo_ = (98 + 8 *·* 100)*/*45 = 19.96.

**Table 2:** Evaluation of variability indices for sorted example time series. CV and SDL use the sample standard deviation; SDL is calculated from ln(*z*) for series without zeros and from ln(*z* + 1) for series containing zeros.

| **y** | CV | SDL | pv<sub>lo</sub> | PV<sub>lo</sub> | PV<sub>hi</sub> |
|---|---|---|---|---|---|
| *y*<sub>1</sub> = (1, 7, 8, 9, 10, 11, 12, 13, 14, 15) | 0.41 | 0.79 | 2.32 | 0.70 | 0.39 |
| *y*<sub>2</sub> = (5, 5, 6, 6, 7, 7, 8, 8, 8, 40) | 1.06 | 0.60 | 1.25 | 0.56 | 0.32 |
| *y*<sub>3</sub> = (5, 5, 8, 8, 10, 10, 12, 12, 15, 15) | 0.36 | 0.40 | 0.67 | 0.40 | 0.34 |
| *y*<sub>4</sub> = (0, 7, 8, 9, 10, 11, 12, 13, 14, 15) | 0.44 | 0.81 | 2.52 | 0.72 | 0.41 |
| *y*<sub>5</sub> = (0, 0, 1, 1, 1, 1, 2, 2, 2, 2) | 0.66 | 0.42 | 0.89 | 0.47 | 0.53 |
| *y*<sub>6</sub> = (0, 0, 1, 3, 10, 12, 15, 17, 20, 22) | 0.85 | 1.26 | 7.23 | 0.88 | 0.70 |
| *y*<sub>7</sub> = (0, 0, 0, 0, 0, 10, 10, 20, 30, 30) | 1.25 | 1.59 | 11.36 | 0.92 | 0.65 |
| *y*<sub>8</sub> = (0, 0, 0, 0, 0, 1, 1, 2, 3, 3) | 1.25 | 0.60 | 1.36 | 0.58 | 0.65 |
| *y*<sub>9</sub> = (0, 0, 0, 0, 0, 0, 0, 0, 1, 99) | 3.13 | 1.45 | 19.96 | 0.95 | 0.38 |
| *y*<sub>10</sub> = (0, 0, 0, 0, 0, 0, 0, 0, 1, 2) | 2.25 | 0.39 | 0.56 | 0.36 | 0.37 |

**Table 3:** Mean *±* SD of the indices across 50 000 simulated time series per regime (upper part) and Spearman correlation with ADM (lower part); numerical values of Figs. 2 and 3.

|  | Normal | Cauchy | Rare | Bimodal |
|---|---|---|---|---|
| *Mean ± SD* |  |  |  |  |
| PV<sub>hi</sub> | 0.151 ± 0.010 | 0.171 ± 0.027 | 0.136 ± 0.004 | 0.505 ± 0.004 |
| PV<sub>lo</sub> | 0.166 ± 0.012 | 0.379 ± 0.177 | 0.540 ± 0.091 | 0.823 ± 0.003 |
| SDL | 0.153 ± 0.012 | 0.364 ± 0.132 | 0.538 ± 0.063 | 1.156 ± 0.010 |
| CV | 0.149 ± 0.011 | 0.947 ± 1.344 | 0.211 ± 0.005 | 0.828 ± 0.004 |
| *Spearman correlation with ADM* |  |  |  |  |
| PV<sub>hi</sub> | 0.94 | 0.60 | 0.87 | 0.14 |
| PV<sub>lo</sub> | 0.91 | 0.70 | 0.48 | 0.72 |
| SDL | 0.86 | 0.71 | 0.52 | 0.73 |
| CV | 0.91 | 0.98 | 0.69 | 0.59 |

A pair (0, *z*) has *d*_hi_ = 1 and *d*_lo_ = *z* (with *z* ≥ 1 for counts). The identity *d*_hi_ = *d*_lo_*/*(1 + *d*_lo_) therefore no longer holds for these pairs: standardized on their own they amount to *z/*(1 + *z*) *<* 1 (only 0.5 for *z* = 1) instead of 1. Without zeros, PV_lo_ ≥ PV_hi_ always holds; with zeros at low abundances this can reverse (*y*_5_, *y*_8_ and *y*_10_ in Table 2), in line with the lower variability that should be indicated if zeros occur at low abundances.

## 3 Results and Comparative Evaluation

### 3.1 Performance on Selected Ecological Abundance Profiles

Table 2 compares CV, SDL, pv_lo_, PV_lo_ and PV_hi_ for ten constructed example series that differ in their distribution and in the number of zero counts. The series are listed in ascending order; since none of the indices depends on the temporal order of the observations, this does not affect the scores.

The CV is dominated by single large values. In *y*_2_, one observation (40) among values of 5–8 produces a CV of 1.06, three times that of the evenly spread *y*_3_ (0.36), and in *y*_9_ one observation (*z*_10_ = 99) produces the highest CV of the table (3.13), although 80% of the observations are zero. The CV also cannot distinguish *y*_7_ from *y*_8_ (both 1.25). The two series differ only in the level of their non-zero abundances (*y*_8_ = *y*_7_*/*10), and because the CV is scale invariant, it does not separate zeros that occur next to abundances of 10–30 from zeros that occur next to abundances of 1–3.

SDL responds to this difference (1.59 for *y*_7_, 0.60 for *y*_8_) and, for example, increases from 1.26 to 1.59 between *y*_6_ and *y*_7_. Its values, however, have no direct meaning on the scale of the abundances: an SDL of 1.59 must be back-transformed to be interpreted, and series with zeros require a constant (here 1) to be added to all abundances before taking logarithms.

The asymmetry of proportional variability becomes apparent when comparing PV_hi_ and PV_lo_. Series *y*_1_ contains a single low value (1) among values of 7–15, series *y*_2_ a single high value (40) among values of 5–8. PV_hi_, PV_lo_ and SDL rate *y*_1_ higher than *y*_2_ (0.39 vs 0.32, 0.70 vs 0.56, 0.79 vs 0.60), whereas the CV rates *y*_2_ higher (1.06 vs 0.41). Because PV_hi_ scales each pair relative to the higher value (Heath, 2006), a single large spike has little influence on it: *y*_2_ obtains 0.32, even slightly less than the evenly spread *y*_3_ (0.34). PV_hi_ cannot, however, reflect how low the lower values are: every pair of a zero and a positive value contributes exactly 1, whether the positive value is 1 or 99. Consequently, *y*_7_ and *y*_8_ obtain the same PV_hi_ (0.65), and *y*_9_ and *y*_10_, which differ only in their largest value (99 vs 2), obtain 0.38 and 0.37. PV_lo_ scales relative to the lower value and therefore grows with the contrast between low and high abundances: it is 0.92 for *y*_7_ and 0.58 for *y*_8_, and 0.95 for *y*_9_ and 0.36 for *y*_10_. The indices also do not jump when a zero appears: replacing the lowest value of *y*_1_ by 0 (series *y*_4_) changes PV_lo_ only from 0.70 to 0.72, PV_hi_ from 0.39 to 0.41, SDL from 0.79 to 0.81 and the CV from 0.41 to 0.44.

The scale of PV_lo_ can be read directly. For *y*_7_, pv_lo_ = 11.36 (PV_lo_ = 0.92): the average of the pairwise differences, each divided by the lower value, is more than eleven (for pairs containing a zero the difference is divided by 1; Section 2.1). Equivalently, PV_lo_*/*(1 − PV_lo_) = pv_lo_; for *y*_7_ this is 0.919*/*0.081 = 11.36 with unrounded values, and 1 + pv_lo_ = 12.36 is the corresponding average ratio of the higher to the lower value (formed after adding 1 to both members of a pair containing a zero).

The context in which zeros occur matters for the interpretation. In count data, the probability of a zero is higher when abundances are low. An index should therefore indicate lower variability if zeros occur in a series of low abundances than if they occur next to high abundances. This can be checked for pairs of series with the same number of zeros: *y*_5_ and *y*_6_ (two zeros), *y*_8_ and *y*_7_ (five) and *y*_10_ and *y*_9_ (eight). PV_lo_ is lower for the low-abundance member of each pair (0.47 vs 0.88; 0.58 vs 0.92; 0.36 vs 0.95), and so is SDL (0.42 vs 1.26; 0.60 vs 1.59; 0.39 vs 1.45). PV_hi_ separates *y*_5_ from *y*_6_ (0.53 vs 0.70), but not *y*_8_ from *y*_7_ (0.65 vs 0.65) or *y*_10_ from *y*_9_ (0.37 vs 0.38), because it scores every pair of a zero and a positive value as 1. SDL achieves its separation only by adding a constant to all abundances, which is known to cause a bias at low abundances regardless of the variance–mean relationship (Gaston & McArdle, 1994). PV_lo_ does not transform the positive abundances, so no constant enters the pairs of positive values; the convention for zero pairs (division by 1) affects only pairs containing a zero, but it refers to the unit of measurement and is therefore meaningful for counts.

In these examples, the CV is driven by single high values and cannot separate scales, PV_hi_ is insensitive to how low the low values are, and PV_lo_ and SDL respond to both, with PV_lo_ on a scale that can be read directly. The next section compares the indices for simulated population dynamics.

### 3.2 Behavior Under Continuous Population Simulations

To evaluate the indices for long time series with different kinds of non-Gaussian dynamics, I simulated population trajectories of 100 time steps under four regimes (Normal, Cauchy, Rare, Bimodal; details in the caption of Fig. 2), with 50 000 simulated series per regime. All abundances are integers of at least 1, so the zero convention of Section 2.1 plays no role here and SDL needs no constant. The indices were computed with the sorting-based algorithm of Section 2.2, which was checked against the pair-wise definition. Each regime probes one property of an index. The Normal regime is a reference in which all measures should agree. The Cauchy regime tests robustness: a few isolated extreme values, one-off booms as well as crashes, should not dominate the index. The Rare regime tests sensitivity: an index should register recurrent crashes of a population that is otherwise stable. The Bimodal regime tests resolution: an index should keep increasing as the gap between two abundance levels widens.

**Figure 2:**
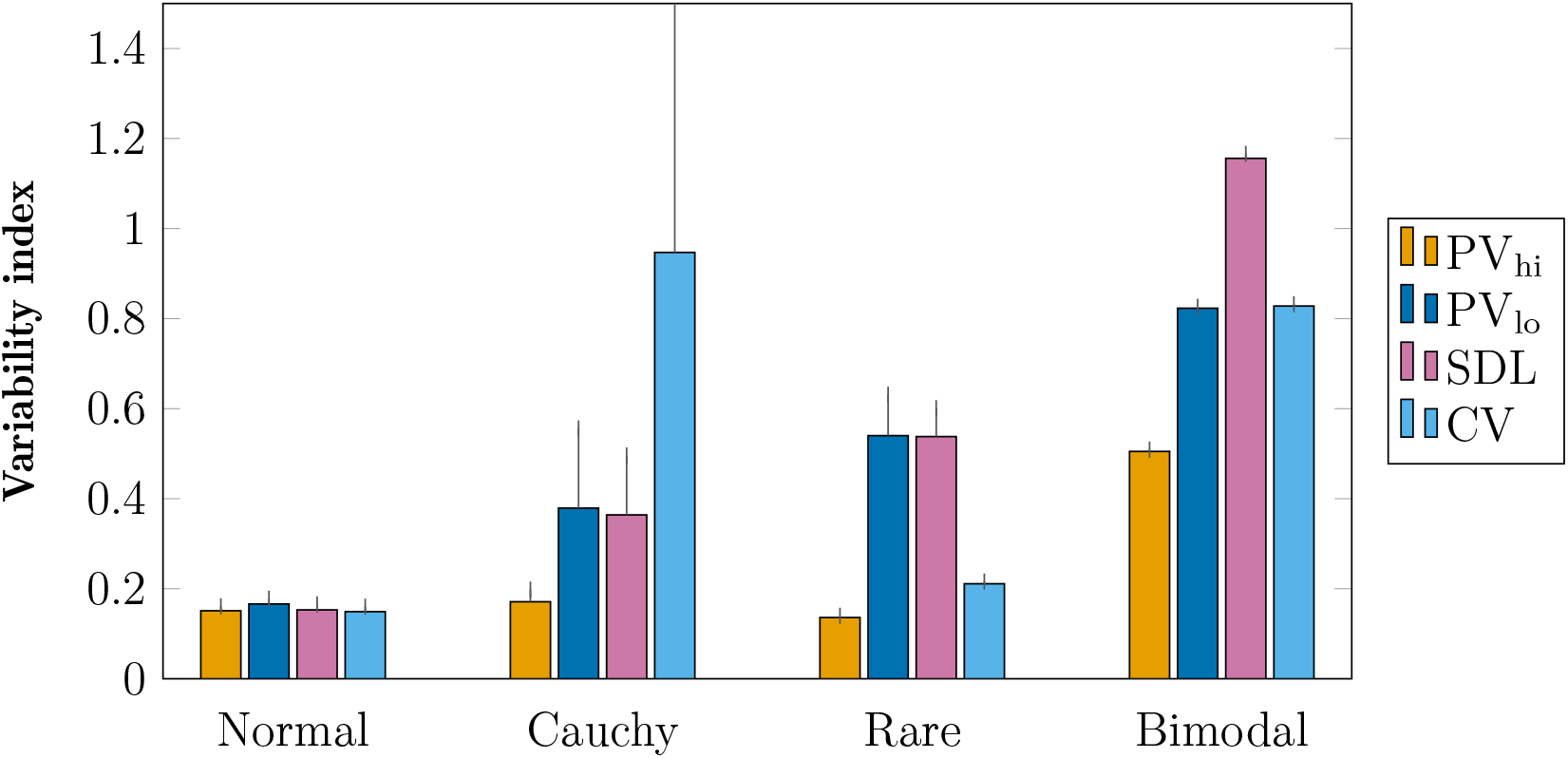
Mean of PV_hi_, PV_lo_, SDL and CV in four simulation regimes (50 000 simulated time series of 100 time steps per regime). Error bars show one standard deviation (SD) above the mean, taken across the simulated series; the upper error bar of CV in the Cauchy regime (mean + SD = 2.29) extends beyond the plotted range. *Normal*: abundances drawn from a Gaussian distribution with mean 1000 and SD 150. *Cauchy*: Cauchy distribution with location 100 and scale 5, rounded to integers, absolute values, lower bound 1 (isolated extreme values in both directions: booms of unbounded size and crashes limited by the lower bound of 1; negative draws are reflected by taking absolute values). *Rare*: 95 abundances from a Gaussian distribution (mean 100, SD 5) and exactly five crashes (5 %) at random positions, with abundances during crashes from a Gaussian distribution (mean 10, SD 5). *Bimodal*: 50 time steps from a Gaussian distribution with mean 100 (SD 10), followed by 50 time steps with mean 10 (SD 1). Gaussian values were rounded to integers; values below 1 were redrawn, so that no series contains zeros.

Fig. 2 shows the mean and the standard deviation (SD) of the indices across the simulated series. Because the indices have different scales, I compare them with their values in the Normal regime, in which they all lie between 0.15 and 0.17. In addition, I calculated Spearman’s rank correlation between each index and the average deviation from the mean, 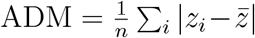, across the simulations (Fig. 3). A high correlation means that an index orders the simulated series like a measure anchored at the mean, and adds little to it. Whether this is a disadvantage depends on the regime. A low correlation is informative only if the index varies systematically between simulations; it also arises when an index is saturated and its remaining variation is mostly noise (see the Bimodal regime below).

**Figure 3:**
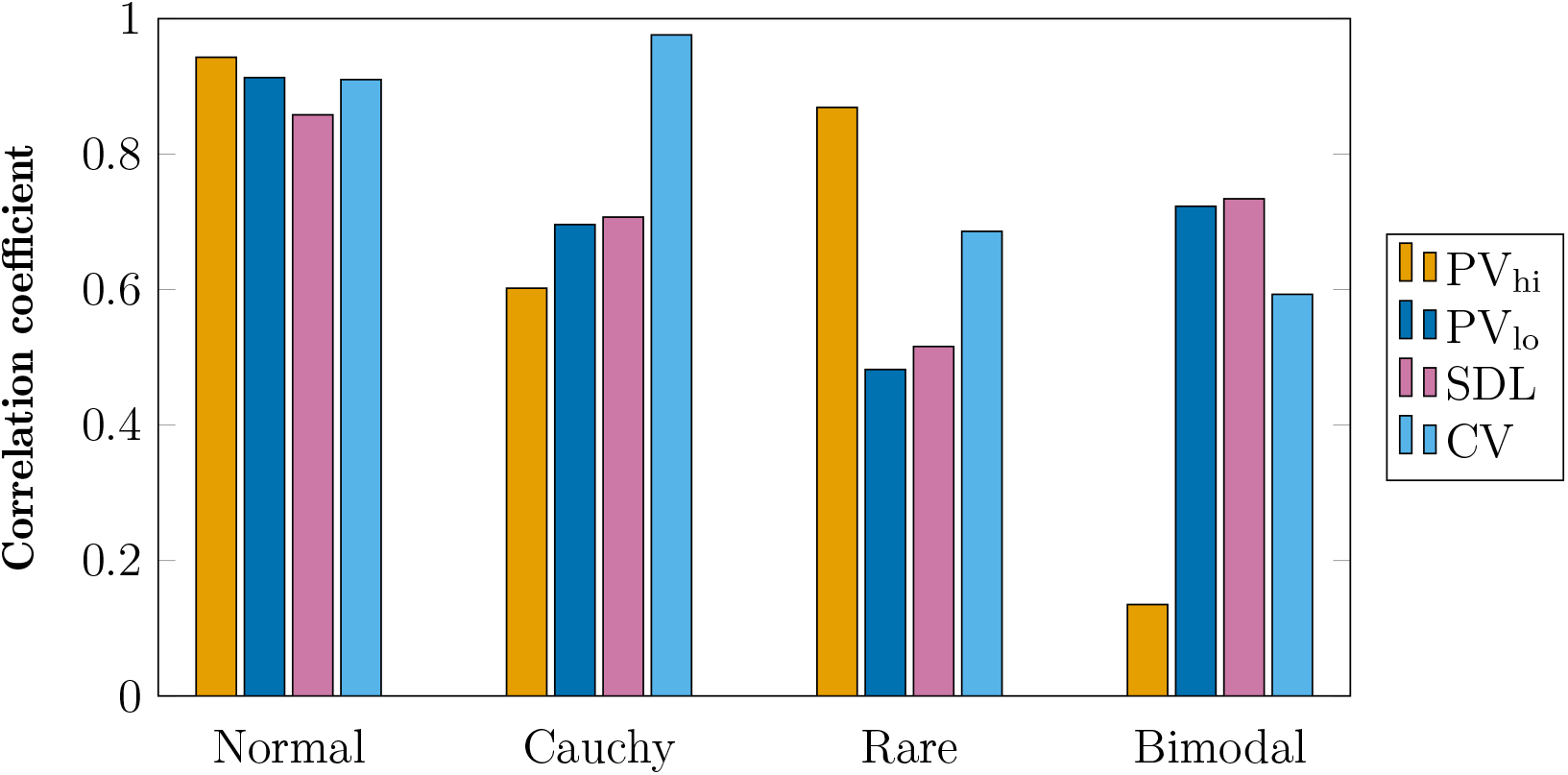
Spearman rank correlation of PV_hi_, PV_lo_, SDL and CV with the average deviation from the mean (ADM), across 50 000 simulated time series of 100 time steps per regime. The correlation of pv_lo_ with ADM is identical to that of PV_lo_, because ranks are unaffected by the standardization. See the caption of Fig. 2 for the four regimes.

#### Normal

The four indices give similar scores (0.15–0.17) and are all strongly correlated with ADM (*r* = 0.86–0.94), as expected for Gaussian data, where all measures reflect the same dispersion.

#### Cauchy

This regime represents isolated extreme values in both directions, that is, one-off population booms and crashes, which are far less frequent than the recurrent crashes of the Rare regime. Taking a tenfold deviation from the location parameter (abundance ≥ 1000 or ≤ 10) as an event, a series contained on average 0.66 events (0.32 booms and 0.34 crashes), about one per 150 time steps compared with one per 20 in the Rare regime. No event occurred in 52% of the series, exactly one in 34% and two or more in 14%; 27% of the series contained at least one boom, 29% at least one crash and 8% both. The two directions are not symmetric: a boom has no upper limit, whereas a crash cannot fall below the lower bound of 1, a hundredfold decline from the location. Here a high score is undesirable, because it shows that a few isolated extreme values determine the result. The CV fails in this respect: it varies strongly between simulations (mean 0.95, SD 1.34) and is almost completely determined by ADM (*r* = 0.98). PV_hi_ stays closest to its Normal level (0.17 *±* 0.03) and is the most robust index here. PV_lo_ (0.38 *±* 0.18) and SDL (0.36 *±* 0.13) react more strongly: their scores are more than twice their Normal values and vary more between simulations, so they are affected by extreme values as well. This is the counterpart of their sensitivity to low abundances (see Rare regime). Compared with the CV, however, this effect is moderate: their correlations with ADM (*r* = 0.70 and 0.71) are clearly lower than that of the CV, i.e. single extremes drive them less. The bounded scale contributes to this moderation: the unstandardized pv_lo_ is as erratic as the CV in this regime (mean 1.34, SD 30.9). Booms and crashes of the same size affect the pairwise indices equally, because *d*_lo_ and the log-ratio underlying SDL depend only on the ratio of two abundances: replacing the booms or the crashes by the location value lowered the mean PV_lo_ by the same amount (0.38 to 0.32 in both cases). Booms matter more in the tail, because only their ratio is unbounded: the 3% of series with a boom of more than a hundredfold account for almost half of the summed pv_lo_ of all series. The CV responds to booms only (mean 2.3 in series with a single boom, 0.40 with a single crash and 0.38 without an event).

#### Rare

This regime represents a conservation-relevant scenario: a population that is stable for 95% of the time but crashes at a frequency of 5%, to about a tenth of its usual abundance. Here a high score is the desired result. Unlike the extremes in the Cauchy regime, the crashes occur regularly and are of moderate depth, so they are exactly the events that an index of demographic instability should register. PV_lo_ (0.54) and SDL (0.54) rise to more than three times their Normal level, whereas the CV (0.21) is only slightly above its Normal value (0.15) and PV_hi_ (0.14) is even slightly lower than in the Normal regime. PV_hi_ thus rates a population that repeatedly loses about 90% of its abundance as no more variable than one with ordinary Gaussian fluctuations of 15%. Whether a larger response also means a better separation from ordinary fluctuations is examined in Section 3.3. Both CV and PV_hi_ are anchored to the high abundances. In PV_hi_ each pair contributes at most 1, and only the stable–crash pairs (475 of 4950, i.e. 9.6%) are affected, each contributing about 0.9. In PV_lo_, such a pair contributes at least about 9 (the ratio of stable to crashed abundance minus one; about 13 on average in the simulation, because crashed abundances scatter and values close to zero weigh disproportionately). This contribution is not capped and grows with the depth of the crash. In Eq. (10), every abundance enters through 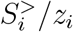, so that the lowest abundances receive the largest weight, whereas in Eq. (9) the weights are bounded by 1.

Concerning ADM, a low correlation is desirable in the Rare regime, because it shows that an index does not merely reproduce the mean-based deviation. PV_hi_ is strongly correlated with ADM (*r* = 0.87) and varies hardly between simulations (SD 0.004): since every series contains exactly five crashes, they add a nearly constant amount to PV_hi_, and the ranking of the simulations mainly follows the same mean-based deviations as ADM. PV_lo_ (*r* = 0.48) and SDL (*r* = 0.52) are less tied to ADM, while the CV lies in between (*r* = 0.69).

Across the simulations, PV_hi_ is strongly correlated with the SD of the stable phase (*ρ* = 0.84), which also drives ADM (*ρ* = 0.66), but only weakly with the mean reciprocal crashed abundance (*ρ* = 0.43). For PV_lo_ the picture is reversed (*ρ* = 0.02 and *ρ* = 1.00, respectively), i.e. it ranks the simulations by the proportional depth of the crashes. As a reference, the stable phase alone (Gaussian, mean 100, SD 5, i.e. 5% relative fluctuation instead of 15% in the Normal regime) yields 0.05 (CV), 0.05 (SDL), 0.06 (PV_lo_) and 0.05 (PV_hi_). Relative to this reference, the five crashes raise the CV about four-fold and PV_hi_ about 2.5-fold, but SDL and PV_lo_ about ten-fold.

#### Bimodal

This regime models a sudden, permanent shift in carrying capacity (50 time steps with a mean of 100 and 50 with a mean of 10). A useful index should keep increasing with the gap between the two levels. PV_hi_ reaches 0.51, which is close to its limit for two equally frequent levels: only the 2500 of 4950 pairs (50.5%) that combine one value from each level contribute substantially (about 0.9 each for a ten-fold gap, at most 1 for larger gaps), and the remaining pairs contribute little. PV_hi_ therefore cannot exceed 0.505 by more than the small contribution of the pairs within each level, however large the gap. PV_lo_ (0.82) is not limited in this way, because pairs are compared with the lower value: its score rises with the ratio between the levels, from 0.82 for a ten-fold to 0.98 for a hundred-fold difference (with the same relative variation within each level). Being bounded, it is compressed close to 1 for larger gaps; for very large contrasts, pv_lo_, which grows in proportion to the ratio, is the more informative scale. In this regime all indices vary very little between simulations (SD ≤ 0.01), so the correlations with ADM reflect sampling noise within the two levels and carry limited information. PV_hi_ is almost uncorrelated with ADM (*r* = 0.14), consistent with a saturated index whose remaining variation carries little information, whereas PV_lo_ and SDL remain correlated with ADM (*r* = 0.72 and 0.73).

In summary, no index is best in all regimes. The CV is dominated by single extreme values under heavy-tailed dynamics and hardly separates the Rare regime from the Normal one. PV_hi_ is the most robust index against heavy-tailed extremes, but it hardly registers rare crashes and levels off when the population switches between two levels. PV_lo_ responded to low abundances in all regimes and gave scores comparable to SDL in the Rare and Cauchy regimes without requiring a transformation of the data. The price of this sensitivity is a stronger response to heavy-tailed extremes (one-off booms as well as crashes) than that of PV_hi_, although far weaker than that of the CV. This makes PV_lo_ the more sensitive index for deep lows, and in these regimes it behaves like SDL. Whether the larger response also means a better separation of crashes from ordinary fluctuations, and how the indices behave in a sustained decline, which is not covered by these regimes, is examined in Section 3.3.

### 3.3 Gradual Decline, Series Length and Separation of Crashes

The regimes of Section 3.2 contain either isolated crashes or a single step between two levels. To examine a sustained decline of the kind that underlies many extinction-risk assessments, I added a fifth regime (Decline). The abundance fluctuates around 100 for the first 50 of 100 time steps and then falls exponentially, i.e. at a constant per-capita rate, to 100*/f* at the last time step. The main case uses *f* = 10, a decline by one order of magnitude as in the IUCN wording quoted in the Introduction; *f* = 100 serves as a sensitivity case. Observations scatter around this path as rounded Gaussian values with a relative SD of 0.15, the level of the Normal regime, and values below 1 were redrawn as before. A stress test with a relative SD of 0.30 is reported as well. Besides the four indices, I included the consecutive disparity index *D* (Fernández-Martínez et al., 2018), the mean absolute log-ratio of consecutive values, as an order-sensitive reference. Each case comprises 50 000 simulated series. Because the indices (except *D*) ignore the temporal order, two questions are of interest: how they respond at the end of the series (Table 4), and when they respond if the series is extended step by step (Table 5). For the second question, an index counted as detecting the decline at a given series length if it exceeded the 95th percentile of the same index in 5000 stable series of the same length and relative SD. This criterion is exploratory: it is one-sided, and no correction for repeated testing along the series was applied. A further question, treated at the end of the section, is whether the stronger response of PV_lo_ and SDL to crashes in the Rare regime means a better separation of crash series from ordinary fluctuations.

**Table 4:** Decline regime: mean *±* SD of the indices at the end of the series (100 time steps) across 50 000 simulated series per case. Stable: abundances fluctuate around 100 throughout; *f* : after 50 time steps the abundance falls exponentially to 100*/f* . Rounded Gaussian scatter with a relative SD of 0.15 (upper part) or 0.30 (lower part). *D* is the consecutive disparity index (Fernández-Martínez et al., 2018).

|  | Stable | *f* = 10 | *f* = 100 |
|---|---|---|---|
| *Relative SD 0.15* |  |  |  |
| PV<sub>hi</sub> | 0.151 ± 0.010 | 0.469 ± 0.006 | 0.604 ± 0.004 |
| PV<sub>lo</sub> | 0.166 ± 0.012 | 0.669 ± 0.009 | 0.932 ± 0.003 |
| SDL | 0.154 ± 0.012 | 0.771 ± 0.016 | 1.530 ± 0.019 |
| CV | 0.150 ± 0.011 | 0.541 ± 0.009 | 0.743 ± 0.007 |
| *D* | 0.173 ± 0.016 | 0.175 ± 0.016 | 0.180 ± 0.018 |
| *Relative SD 0.30* |  |  |  |
| PV<sub>hi</sub> | 0.279 ± 0.019 | 0.508 ± 0.010 | 0.635 ± 0.007 |
| PV<sub>lo</sub> | 0.358 ± 0.056 | 0.709 ± 0.024 | 0.935 ± 0.005 |
| SDL | 0.350 ± 0.050 | 0.833 ± 0.038 | 1.547 ± 0.033 |
| CV | 0.299 ± 0.023 | 0.614 ± 0.020 | 0.808 ± 0.018 |
| *D* | 0.372 ± 0.043 | 0.373 ± 0.042 | 0.368 ± 0.042 |

**Table 5:** Decline regime (*f* = 10), expanding window: share of decline series in which the index exceeds the 95th percentile of the same index in stable series of the same length, +*d* time steps after the onset of the decline (2000 decline series, 5000 stable reference series; relative SD 0.15 in the upper and 0.30 in the lower part). The nominal share for a stable series is 0.05.

|  | +10 | +20 | +30 |
|---|---|---|---|
| *Relative SD 0.15* |  |  |  |
| PV<sub>hi</sub> | 0.64 | 1.00 | 1.00 |
| PV<sub>lo</sub> | 0.66 | 1.00 | 1.00 |
| SDL | 0.67 | 1.00 | 1.00 |
| CV | 0.63 | 1.00 | 1.00 |
| *D* | 0.06 | 0.07 | 0.07 |
| *Relative SD 0.30* |  |  |  |
| PV<sub>hi</sub> | 0.13 | 0.85 | 1.00 |
| PV<sub>lo</sub> | 0.06 | 0.27 | 0.99 |
| SDL | 0.07 | 0.32 | 1.00 |
| CV | 0.14 | 0.85 | 1.00 |
| *D* | 0.06 | 0.06 | 0.06 |

All order-blind indices responded clearly to the decline (Table 4). For *f* = 10 and a relative SD of 0.15, PV_hi_ rose from 0.15 to 0.47 (3.1-fold), PV_lo_ from 0.17 to 0.67 (4.0-fold), CV 3.6-fold and SDL 5.0-fold. In contrast to the Rare regime, where it did not register the crashes, PV_hi_ is thus clearly sensitive to a sustained decline: a decline places many observations below earlier ones, and every one of the many resulting pairs contributes. The advantage of PV_lo_ over PV_hi_ is therefore moderate (response ratio 1.3 for *f* = 10 and 1.4 for *f* = 100); PV_lo_ lies close to the CV, and SDL responds most. *D* did not respond (0.173 in the stable series and 0.175 at *f* = 10), as expected for an index of consecutive changes, which remain small in a slow decline.

The indices differ in how they register the depth of the decline. From *f* = 10 to *f* = 100, PV_hi_ increased by 0.14 (0.47 to 0.60), whereas PV_lo_ increased from 0.67 to 0.93 and the unstandardized pv_lo_ from 2.0 to 13.8, so that the contrast between the two depths remains visible on its scale. For rank discrimination, however, the advantage is small: at both levels of scatter, all four indices separated *f* = 10 from *f* = 100 perfectly (AUC 1.00), and at a relative SD of 0.30 only the CV lost some resolution between *f* = 50 and *f* = 100 (AUC 0.95, against 0.995 for PV_hi_ and 1.00 for PV_lo_ and SDL).

In the expanding window, the indices stayed at their stable level until the decline began and then rose as the series extended into it. For *f* = 10 and a relative SD of 0.15, the mean PV_lo_ was 0.17 at 30 and at 50 time steps, 0.20 at 60, 0.31 at 70 and 0.67 at 100 (Fig. 4a). This is the dependence on series length discussed in Section 3.2 and in the Discussion, but in a non-stationary setting: it reflects the decline and not a bias of the index, and in stable Normal series PV_lo_ stayed at 0.17 for all lengths. At a relative SD of 0.15, the four indices detected the decline equally fast (Table 5): ten time steps after its onset, the share of detections was 0.66 for PV_lo_, 0.64 for PV_hi_, 0.67 for SDL and 0.63 for the CV, and after twenty steps it was 1.00 for all four. *D* stayed at the nominal level (0.06 to 0.07).

**Figure 4:**
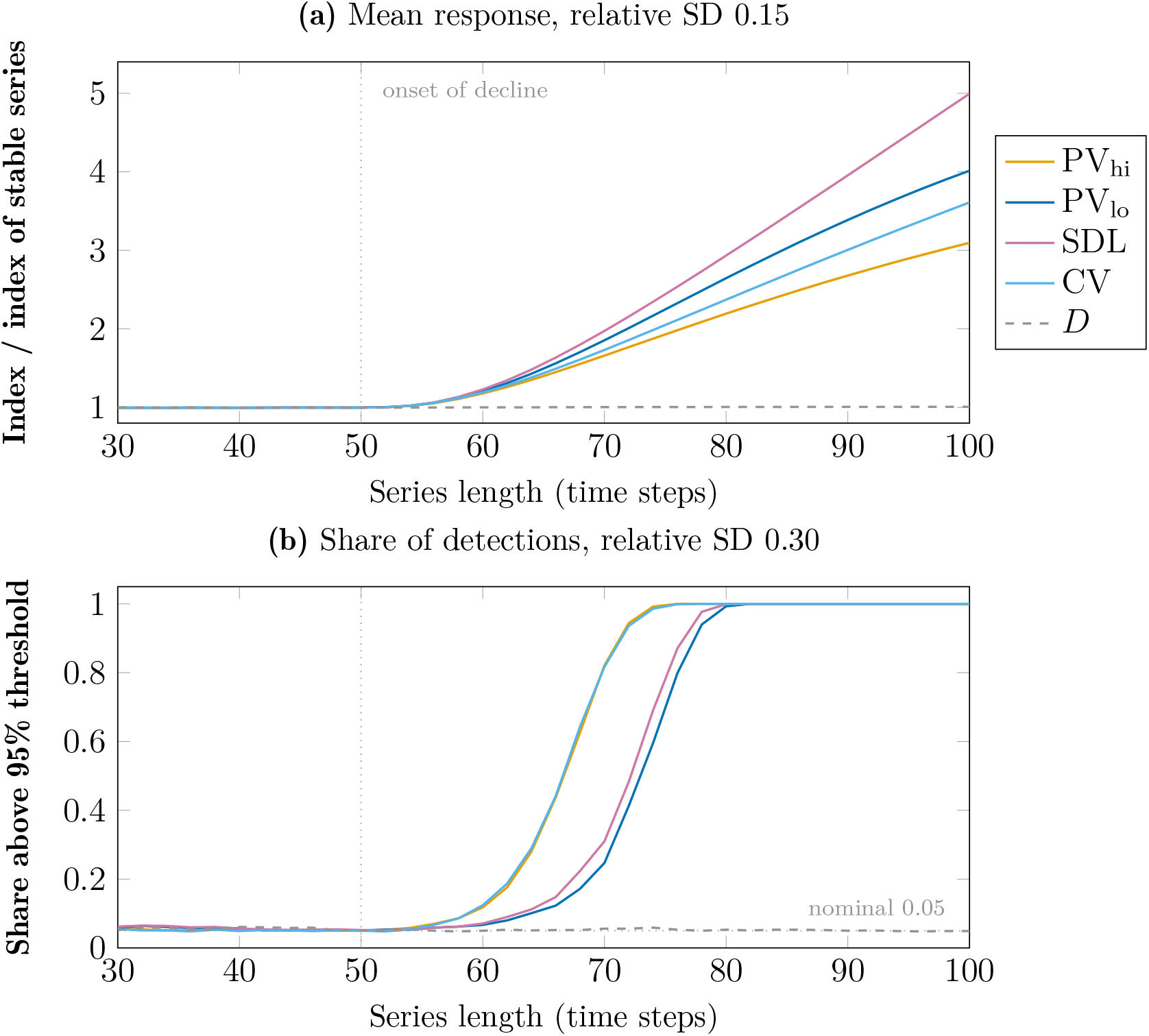
Decline regime (*f* = 10; the decline begins after 50 time steps), expanding window. (a) Mean index at each series length divided by the mean of the same index in stable series of that length; relative SD 0.15. (b) Share of decline series in which the index exceeds the 95th percentile of the same index in stable series of the same length; relative SD 0.30. The dotted vertical line marks the onset of the decline, the dotted horizontal line in (b) the nominal share of 0.05. PV_hi_ and CV almost coincide in (b). 2000 decline series and 5000 stable reference series per panel.

The stress test shows the price of the weight that PV_lo_ puts on low values. At a relative SD of 0.30, PV_lo_ and SDL detected the decline later than PV_hi_ and the CV (Fig. 4b): twenty steps after the onset the shares were 0.27 and 0.32 against 0.85 for both PV_hi_ and CV, and thirty steps after the onset PV_lo_ reached 0.99. For *f* = 100 the shares after ten steps were 0.14 (PV_lo_) and 0.18 (SDL) against 0.50 (PV_hi_) and 0.49 (CV), and all four indices reached 1.00 after twenty steps. The reason is that scatter at this level produces low values even in stable series, so that the distribution of PV_lo_ in the stable phase is wider (SD 0.056 at a mean of 0.36, against 0.019 at 0.28 for PV_hi_) and the detection threshold higher.

Is the stronger response of PV_lo_ and SDL to crashes in the Rare regime (Section 3.2) an advantage in telling crashes from ordinary fluctuations? A larger response alone does not show this, because it concerns the scale of an index and not its ability to separate two kinds of series. I therefore compared series with five crashes at random positions (to 100*/f*, with a relative SD of 0.5 of the crashed abundance, as in the Rare regime) with series of ordinary Gaussian fluctuations around 100, using the AUC, i.e. the probability that a crash series has a higher index value than an ordinary series. The AUC depends only on ranks, so that a monotone standardization such as Eq. (5) does not change it. Crash and ordinary series share the same relative SD of the fluctuations. It was either fixed, which corresponds to a known baseline, or drawn at random for every series, which corresponds to a comparison of populations whose ordinary variability is unknown.

If the baseline was known, the indices separated crashes equally well (Table 6). At a relative SD of 0.05 the AUC was 1.00 for all indices and crash depths, at 0.15 it was 1.00 for all indices from *f* = 5 on, and at 0.30 the five indices were within 0.03 of each other. The picture changed if the baseline was unknown. For a relative SD between 0.05 and 0.30 and *f* = 10, the AUC was 0.77 for PV_hi_ and 0.86 for the CV, but 1.00 for PV_lo_ and SDL; for *f* = 2 it was 0.63 and 0.66 against 0.75 and 0.79. For a relative SD between 0.02 and 0.50 all indices separated less well, but the order remained (0.66 and 0.69 against 0.91 at *f* = 10). The consecutive disparity index *D*, which responds to the large consecutive changes that crashes produce, lay between the two groups (0.94 at *f* = 10 for the narrower range). A plausible reason is that PV_hi_ and the CV vary roughly in proportion to the amplitude of ordinary fluctuations, so that their values for ordinary series of different amplitude overlap with those of crash series, whereas a crash raises PV_lo_ and SDL about ten-fold (Section 3.2), more than a change of the amplitude does.

**Table 6:** Separation of series with five crashes from series of ordinary Gaussian fluctuations, measured by the AUC (probability that a crash series has a higher index value than an ordinary series; 0.5: no separation, 1: complete separation). Crash series: Gaussian fluctuations around 100 plus five crashes at random positions to 100*/f* (relative SD of the crashed abundance 0.5). Both kinds of series share the same relative SD of the fluctuations, which was either fixed (known baseline) or drawn uniformly at random for every series (unknown baseline; ranges in brackets). 5000 series per class and case; *D* is the consecutive disparity index.

| *f* | PV<sub>hi</sub> | PV<sub>lo</sub> | SDL | CV | *D* |
|---|---|---|---|---|---|
| *Known: relative SD 0.15* |  |  |  |  |  |
| 2 | 0.974 | 0.990 | 0.994 | 0.984 | 0.972 |
| 5 | 1.000 | 1.000 | 1.000 | 1.000 | 1.000 |
| 10 | 1.000 | 1.000 | 1.000 | 1.000 | 1.000 |
| *Known: relative SD 0.30* |  |  |  |  |  |
| 2 | 0.773 | 0.766 | 0.769 | 0.768 | 0.752 |
| 5 | 0.967 | 0.966 | 0.978 | 0.964 | 0.973 |
| 10 | 0.989 | 0.991 | 0.999 | 0.990 | 0.997 |
| *Unknown: relative SD in* [0.05, 0.30] |  |  |  |  |  |
| 2 | 0.625 | 0.746 | 0.788 | 0.659 | 0.668 |
| 5 | 0.730 | 0.972 | 0.986 | 0.807 | 0.854 |
| 10 | 0.766 | 0.998 | 0.999 | 0.862 | 0.944 |
| *Unknown: relative SD in* [0.02, 0.50] |  |  |  |  |  |
| 2 | 0.557 | 0.610 | 0.625 | 0.572 | 0.573 |
| 5 | 0.633 | 0.783 | 0.796 | 0.657 | 0.690 |
| 10 | 0.661 | 0.907 | 0.912 | 0.692 | 0.768 |

The advantage of PV_lo_ and SDL over PV_hi_ and the CV in telling crashes from ordinary fluctuations thus exists only if the ordinary variability of the population is unknown, and it concerns isolated deep lows. With a known baseline the indices are equivalent, and SDL separated as well as or slightly better than PV_lo_ in all cases of Table 6. SDL is not uniformly better, however. In additional runs beyond the range of the table (one to fifteen crashes, relative SD up to 0.50, crash depth up to *f* = 100; not shown), SDL separated better when several crashes occurred, whereas PV_lo_ was clearly better for a single crash of the maximal depth of the Cauchy regime (*f* = 100, i.e. to about 1) in strongly scattered series (AUC 0.92 versus 0.86 at a relative SD of 0.50 with known baseline, and 0.96 versus 0.85 for an unknown SD between 0.02 and 0.50; at a relative SD of 0.30 or less the difference was at most 0.004). Unlike the extremes of the Cauchy regime, these are collapses only, the events that matter for extinction risk, but a collapse to a hundredth of the usual abundance is conspicuous without an index, so this advantage should not be overrated. The advantage did not carry over to the decline regime: with a baseline of unknown amplitude, the four indices separated decline series from stable series of the same length with AUC values within 0.04 of each other (series of 70 and 100 time steps, *f* = 2 and *f* = 10; not shown). With a known baseline, PV_lo_ and SDL differed by less than 0.011 in either direction, with PV_lo_ marginally better for the shallow decline (*f* = 2; not shown).

In sum, the indices differ less in a sustained decline than under isolated crashes, and the differences depend on what is known about the population. All order-blind indices respond to a decline, PV_hi_ included; the distinctive contribution of PV_lo_ there is the scale on which the depth of the decline is expressed, not earlier detection, and in strongly scattered series it detects later than PV_hi_ and the CV. In telling crashes from ordinary fluctuations, PV_lo_ and SDL were superior to PV_hi_ and the CV only if the ordinary variability was unknown, which is the situation when many populations are screened, and PV_lo_ did not outperform SDL by a noticeable margin in any of these comparisons, except for a single crash of maximal depth in strongly scattered series. A possible interpretation, which these simulations do not test separately, is that the advantage arises for isolated deep lows, which are not a feature of ordinary fluctuations, whereas a decline shifts the bulk of the observations and then puts the CV and PV_hi_ at no disadvantage. As the other order-blind indices, PV_lo_ cannot tell a directional decline from recurrent crashes, and *D*, which is order-sensitive, does not respond to a gradual decline at all, so that the two kinds of index answer different questions.

## 4 Discussion

Which feature of a time series a variability index is anchored to determines which populations it ranks as unstable. The risk of extinction is governed less by how high abundances peak than by how far they fall, because demographic stochasticity and random catastrophes act mainly on small populations (Pimm, 1991; Lande, 1993). Moreover, population change is naturally expressed as a ratio: a population that falls from 1000 to 100 individuals has lost the same proportion as one that falls from 100 to 10. Conservation thresholds follow the same logic. The IUCN criteria, for example, refer to fluctuations of more than one order of magnitude (IUCN Standards and Petitions Committee, 2024). An index that is anchored to lower abundances and can be read as an average ratio is therefore well suited to ask how deep the lows of a population are.

The comparisons show that the three proportional indices register different features. PV_hi_ is the most robust against heavy-tailed extremes but is largely blind to the depth of lows; PV_lo_ and SDL respond strongly to deep lows. That PV_lo_ resembles SDL is expected, because SDL also aggregates all pairwise ratios, only as a squared logarithm instead of a ratio. The more sensitive response translated into better separation of crashes from ordinary fluctuations only when the ordinary variability of a population was unknown and the lows were isolated, which is the situation when many populations are screened; with a known baseline, or during a sustained decline, the indices differed little. PV_lo_ should therefore not be understood as a more powerful substitute for SDL. Its contribution lies in the combination of properties: to my knowledge, no established measure unites sensitivity to the depth of lows, a scale that can be read directly in abundance ratios, the inclusion of zeros without a constant added to all observations, and a computation that remains feasible for long series. This combination is not necessarily the best possible one, but it fills a gap between PV_hi_ and SDL.

Following Heath (2006), who stressed that proportional variability is not an exclusive alternative to other approaches but most informative in conjunction with them, I suggest using PV_lo_ as a complement to other measures rather than as a replacement. Measures such as the CV or PV_hi_, which respond differently to extreme values, are natural companions. Like all indices considered here, PV_lo_ ignores the temporal order. Trends raise it, as Fernández-Martínez et al. (2018) showed for PV_hi_, and it cannot distinguish a directional decline from recurrent crashes. Where chronology matters, it should be combined with a measure based on consecutive observations, such as the consecutive disparity index *D* (Fernández-Martínez et al., 2018).

Several further limitations apply. The standardization makes PV_lo_ depend on series length when rare events occur, in contrast to PV_hi_ (variance exponent under rare crashes 0.37, compared with 0.02 for PV_hi_ and 0.30 for SDL), but the unstandardized pv_lo_ does not, so that reporting both allows the effect to be checked. The zero convention refers to the unit of measurement and is natural for counts of individuals but must be chosen deliberately for biomass or density. Because low abundances carry the most weight, PV_lo_ is also the index most affected by observation error where counts are least reliable. Finally, no empirical time series were analyzed, so the relevance of PV_lo_ for persistence and extinction risk is argued from its properties and remains to be tested on monitoring data.

The efficient algorithm makes it possible to apply PV_lo_ to large monitoring databases and to screen thousands of populations for deep lows, a task for which a ratio-scale index without data transformation offers a practical alternative to SDL.

## Code availability

Jupyter Notebooks with code used for analyses are available at

https://github.com/FT81/prop-var

## Notes

https://github.com/FT81/prop-var

### Competing Interest Statement

The authors have declared no competing interest.

### Summary of Updates

Changes relative to the previous version (v3). This version is substantially revised. (1) The combined index PV (the mean of the standardized indices relative to lower and higher abundances) has been dropped. The paper now concentrates on PV_lo, the standardized average proportional difference relative to lower abundances, and compares it with Heath's index (PV_hi), CV and SDL. The relation of PV_lo to PV_hi and its interpretation as an average abundance ratio are derived. (2) Pairs containing a zero are now divided by one individual instead of being scaled by e/2, so scores for series with zeros differ from the previous version. (3) The sorting-based algorithm is derived in full, including ties and zero counts. (4) The simulations are described in detail and extended by an analysis of series-length dependence, a gradual-decline regime with an expanding window, the consecutive disparity index D as an order-sensitive reference, and a comparison of crash series with ordinary fluctuations. (5) The conclusions are more cautious: PV_lo is more sensitive to deep lows than PV_hi and CV and comparable to SDL, but it separates crashes from ordinary fluctuations better than PV_hi and CV only if the ordinary variability is unknown. It does not detect gradual declines better, PV_hi is more robust against heavy-tailed extremes, and the series-length dependence is similar to that of SDL. The statement in v3 that CV and PV_hi are unsuitable for detecting long-term decline is not retained. (6) The consecutive index CD proposed in v3 is no longer part of this paper. (7) The title and abstract have been revised; Figures 1-4 and Tables 3-6 have been updated or newly added; the references have been updated; and the code link now points to the prop-var repository.

